# Dissemination of *Cryptococcus neoformans* via localised proliferation and blockage of blood vessels

**DOI:** 10.1101/184200

**Authors:** Josie F Gibson, Robert J Evans, Aleksandra Bojarczuk, Richard Hotham, Anne K Lagendijk, Benjamin M Hogan, Philip W Ingham, Stephen A Renshaw, Simon A Johnston

**Affiliations:** Department of Infection, Immunity and Cardiovascular disease, Medical School, University of Sheffield, UK.; The Bateson Centre, University of Sheffield, Sheffield, UK; Institute of Molecular and Cell Biology, Agency of Science, Technology and Research (A-Star), Singapore; Division of Genomics of Development and Disease, Institute for Molecular Bioscience, University of Queensland, Brisbane, Australia; Lee Kong Chian School of Medicine, Nanyang Technological University, Singapore.

## Abstract

*Cryptococcus neoformans* is an opportunistic fungal pathogen that can cause life-threatening cryptoccocal meningitis, predominantly within immunocompromised individuals. Cortical infarcts are observed in as many as 30% of cryptococcal meningitis cases, being particularly common in severe infection. Limited clinical case studies suggest infarcts are secondary to vasculitis and blood vessel damage caused by cryptococcal infection. However, the cause of infarcts in cryptococcal infection has not been determined. To examine potential causes of vascular damage and cryptococcal dissemination in cryptococcal infection, the zebrafish *C. neoformans* infection model was used. We demonstrate that spread of cryptococci from the vasculature occurs at sites where cryptococci grow within the blood vessels, originating from a single or small number of cryptococci. We find that cryptococcal cells become trapped within the vasculature and can proliferate there resulting in vasodilation. Localised cryptococcal growth in the vasculature is also associated with sites of dissemination – in some cases simultaneously with a loss of blood vessel integrity. Using a cell-cell junction protein reporter (VE-cadherin) we identified sites dissemination associated with both intact blood vessels and where vessel rupture occurred. Thus, we have identified a mechanism for blood vessel damage during cryptococcal infection that may represent a cause of the vascular damage and cortical infarction observed in cryptococcal meningitis.

**Author summary:** Human infection by the fungal pathogen, *Cryptococcus neoformans*, can lead to life-threatening cryptococcal meningitis. In severe cases of cryptococcal meningitis, a lack of blood supply can cause tissue death and a resulting area of dead tissue (infarct) in the brain. Although vasculature inflammation in known to occur in cryptococcal meningitis, the cause of infarcts in unknown. Using a zebrafish model of cryptococcal infection, the growth and dissemination of fungal cells was observed over time. We show that cryptococcal cells become trapped and proliferate in the vasculature, resulting in cryptococcoma that damage the blood vessels. We propose that vessel damage results from increased blood pressure caused by cryptococci blocking blood vessels suggesting that the vascular damage that ensues on cryptococcoma formation may in turn be a cause of infarct formation seen in cryptococcal meningitis.

## Introduction

*Cryptococcus neoformans* is an opportunistic fungal pathogen causing infection primarily in immunocompromised patients. Dissemination of infection to the central nervous system (CNS) results in life threatening cryptococcal meningitis and encephalitis. *C. neoformans* is a significant pathogen of HIV/AIDs positive individuals, with mortality rates as high as 70% in sub-Sharan Africa and with cryptococcal meningitis ultimately responsible for 15% of all AIDS related deaths worldwide (1).

Primary infection occurs through environmental exposure to inhaled cryptococcal spores. *C. neoformans* is first encountered by alveolar macrophages, leading to an immune response where infection is resolved through granuloma formation in healthy individuals (2). However, immunocompromised individuals cannot control initial infection, with pulmonary cryptococcal infection preceding dissemination and invasion of the brain (3–5). Cryptococcal cells proliferate within the initial infection site in the lung before escaping and disseminating infection to multiple organs, most significantly the CNS.

The mechanism of *C. neoformans* entry into the CNS is unknown, but must involve breach of the protective barriers of the brain, for example the blood-brain barrier (BBB). An active cell-crossing mechanism has been described *in vitro*, in which cryptococcal cells are able to transcytose through microvascular endothelial cells (6). In addition, *C. neoformans* may be able to cross the BBB between tight junctions, based on evidence of damage to the major tight-junction transmembrane protein occludin after incubation with cryptococcal cells (Chen et al., 2003). On the other hand, host responses may also enable cryptococcal dissemination into the brain. *In vivo* studies have suggested that cryptococcal cells use host macrophages as Trojan horses to cross the BBB, (7). Whilst various factors may play a role in *C. neoformans* ability to cross the blood brain barrier, each method requires the presence of cryptococcal cells within brain blood vessels.

A small number of clinical studies have suggested that blood vessel damage and bursting may also facilitate cryptococcal dissemination, however the mechanism of blood vessel damage is not known. Case reports indicate that cortical infarcts are secondary to cryptococcal meningitis, and suggest a mechanism whereby resulting inflammation may lead to damage to blood vessels (8–10). This is supported by the observation that cerebral infarcts are observed in meningitis caused by either cryptococcal or tuberculosis infection (11). Alternatively, toxins or physical damage from pathogens may directly cause vascular inflammation. In retrospective studies of human cryptococcal infection, instances of vascular events resulting in infarcts were seen in 30.3% of cases, predominantly within severe cases of cryptococcal meningitis (12). The non-trivial incidence rate of infarcts, in addition to their associated higher mortality risk, makes the investigation of blood vessel damage in cryptococcal infection an important research question, specifically with respect to the identification of potential therapeutic targets or the modification of clinical management of patients. Furthermore, cryptococcal cells can become physically trapped in mouse brain blood vessels (13). This is consistent with clinical post-mortem reports showing cryptococcal cells invading the brain, observed both in the perivascular spaces and located next to brain capillaries that have cryptococcal masses or cryptococcoma present (14,15), suggesting that cryptococcal proliferation within the brain blood vessels may lead to vessel damage and subsequent invasion of the brain.

Long term *in vivo* analysis of cryptococcal infection is not possible in rodent or leporine models; the ease of imaging cryptococcal infection in zebrafish, by contrast, enables visualisation of infection dynamics, with many cryptococcal infection characteristics, including brain dissemination, recapitulated in this model (16,17). Furthermore, the cryptococcal zebrafish model enables high quality imaging of host pathogen interactions, including throughout the vasculature, that are of specific relevance for this study. Notably, a high fungal burden within a particular zebrafish cranial blood vessel, is correlated with a higher chance of tissue invasion into the brain (Tenor et al., 2015). Based on this finding together with the observation that cryptococcal cells can get trapped in small blood vessels in the brain, we decided to investigate the role of cryptococcal expansion within blood vessels as a route of dissemination. We postulated that once trapped within a brain blood vessel, *C. neoformans* continues to proliferate, leading to physical damage of the vasculature and eventual dissemination and invasion of the surrounding tissue.

In this study we observed cryptococcal cells becoming trapped and then proliferating within the vasculature in a manner similar to that seen in murine models. Analysis of the dynamics of infection, via mixed infection of two fluorescent strains of *C. neoformans*, demonstrated that cryptococcomas within small blood vessels were responsible for overwhelming systemic infection. Localised expansion of *C. neoformans* was observed at sites of dissemination into surrounding tissue. Using a new VE-cadherin transgenic reporter line, we identified physical damage to the vasculature at sites of cryptococcal colonisation and found that blood vessels respond to their colonisation via expansion. Taken together, our data demonstrate a previously uncharacterised role of cryptococcal proliferation as a physical dissemination route from the vasculature.

## Results

### Individual cryptococcal cells trapped in blood vessel result in cryptococcomas in the vasculature

Infection of zebrafish with a low dose of ~25cfu directly into the bloodstream of *C. neoformans* resulted in single cryptococcal cells trapped in the vasculature. Exploiting the unique capacity of zebrafish for long term, non-invasive in vivo imaging, we found that the sites of single or very small numbers of trapped cells progressed to form cryptococcal masses or cryptococcomas within blood vessels (Fig 1). These data suggest that localised clonal expansion results in cryptococcoma formation. However, with this approach we could not determine whether cryptococcomas formed via the clonal expansion of individual cryptococci or an accumulation of cells at a single site.

**Figure 1.**
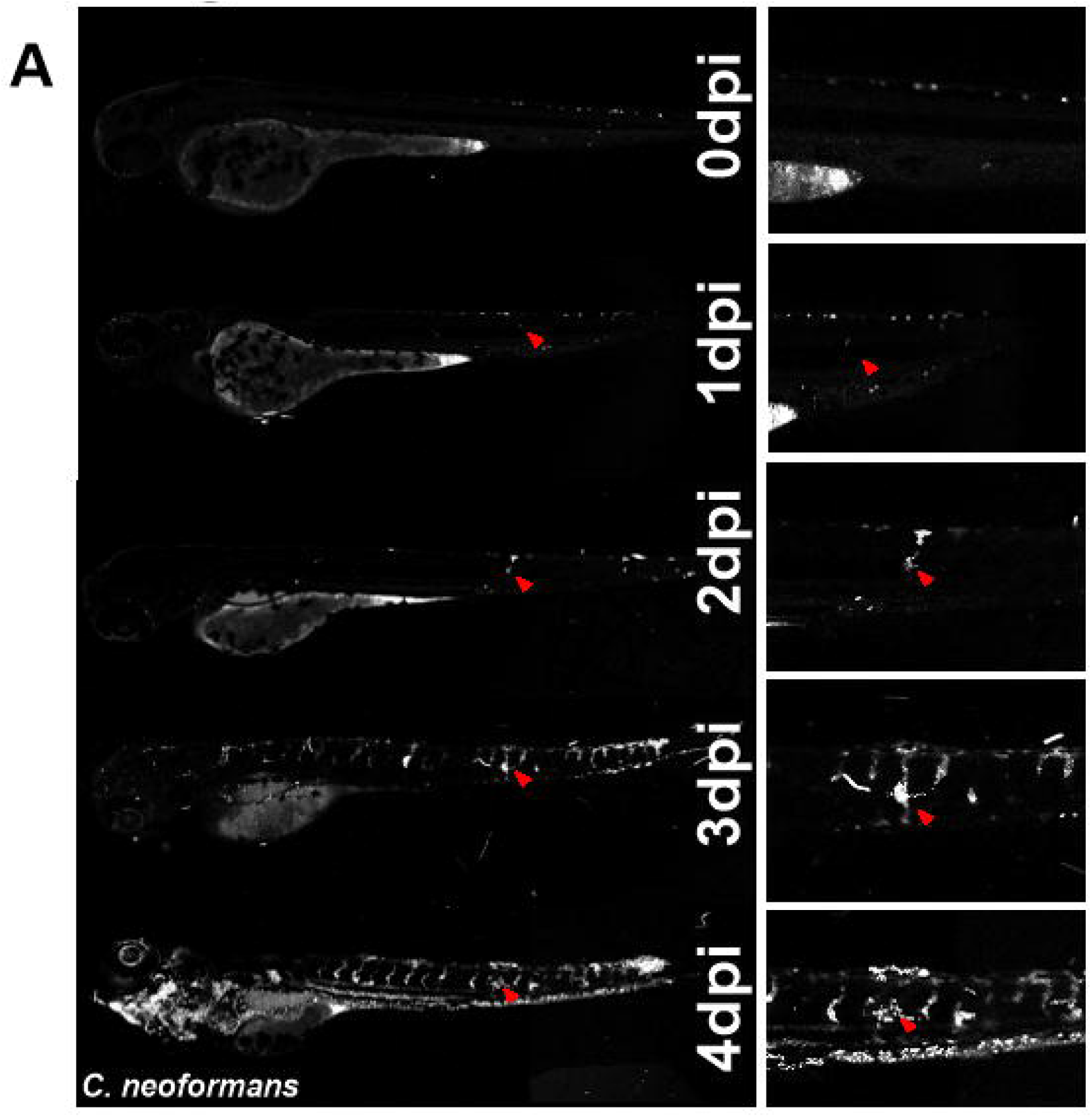
Dissemination occurs at sites of clonal expansion. **A.** Infection of 2dpf AB larvae with 25cfu of a 5:1 ratio of GFP:mCherry KN99 *C. neoformans*. Larvae were imaged until 8dpf, or death (n=3, in each repeat 7, 10 and 12 larvae were used). In this case an mCherry majority overwhelming infection was reached. Infection progression from 0dpi (day of infection imaged 2hpi), until 4dpi. Red arrows follow the formation of an individual cryptococcoma and its dissemination.

### Clonal expansion of individual cryptococci does not correlate with high fungal burden

Several bacterial pathogens have been demonstrated to establish disease via clonal expansion of an individual or small number of pathogens through a population “bottle-neck”. However, we hypothesised that this was not the case for cryptococcosis where there is uncontrolled infection, as it is the growth of extracellular yeast that has been observed during infection of immunocompromised hosts (4,5,16,18). Initially, we injected a 1:1 ratio of GFP and mCherry-labelled cryptococci and found that single colour infections were rare. Therefore, we decided to use a skewed ratio so that we could better quantify the likely hood of a population “bottleneck” during the progression of cryptococcal infection. We injected a 5:1 ratio of GFP and mCherry-labelled cryptococci and followed the infections for up to 7dpi (days post infection). In 51.6% of all infected larvae, a high fungal burden end-point was demonstrated, with either cryptococci predominantly GFP positive, predominantly mCherry positive cryptococci, or a mixed outcome of both GFP and mCherry positive cryptococci (Fig. 2A). Interestingly, a mixed final outcome group was not a rare occurrence (Fig. 2B). The high proportion of mixed GFP and mCherry overwhelming infections demonstrated that a single cryptococcal cell was likely not to give rise to the entire overwhelming infection, when compared to single colour outcomes, which would be expected if a single cryptococcal cell was responsible (Fisher’s exact test p<0.0177). The predominantly GFP positive outcome group was observed most often, but only 56.25% of all endpoints, lower than expected given the initial 5:1 ratio of differentially labeled cells injected. To determine whether the injected ratio was responsible for overwhelming infection we measured the exact initial inoculum ratio. While a 5:1 ratio of GFP:mCherry was injected into each larvae, the actual number and ratio of cryptococcal cells varied between individual fish (Fig. 2C, SFig 1). When compared to the final infection outcome, the initial infection inoculum ratio was not significantly different between GFP and mixed outcomes (Fig. 2D). Furthermore, there was no relationship shown between the initial ratio and final outcome ratio (Fig. 2E), confirming our conclusion that inoculum ratio did not directly determine infection outcome. An example of expected results had inoculum ratio been responsible for final ratio is shown in (Fig. 2F).

**Figure 2.**
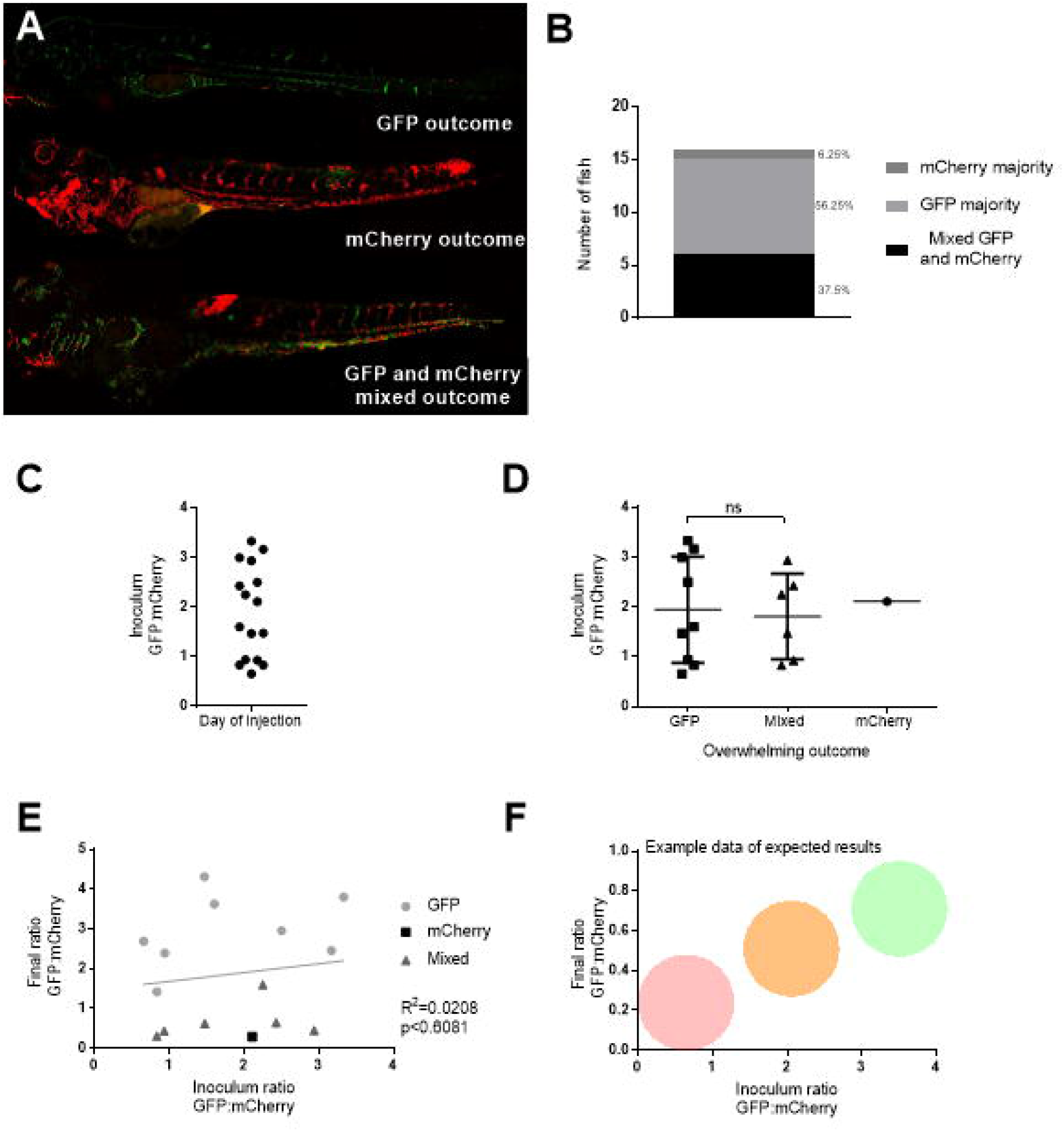
Initial inoculum does not determine infection outcome. Infection of 2dpf AB larvae with 25cfu of a 5:1 ratio of GFP:mCherry KN99 *C. neoformans*. Larvae were imaged until 8dpf, or death (n=3, in each repeat 7, 10 and 12 larvae were used) **A.** GFP majority infection outcome, mCherry infection outcome or a Mixed GFP and mCherry infection outcome (n=3, 16 larvae) **B.** Proportion of each overwhelming infection outcome observed, GFP, mCherry or mixed **C.** Actual injected ratios of GFP:mCherry at 2hpi **D.** Actual injected GFP:mCherry ratios for each overwhelming outcome (n=3, +/- SEM, Man-Whitney t-test ns=not significant) **E.** Inoculum ratio of GFP:mCherry, against final GFP:mCherry ratio at overwhelming infection stage (Linear regression R^2^=0.0208, p<0.6081, n=3, 16 larvae) **F.** A sample data set of expected results representing how graph E may appear if initial inoculum did control final outcome

### Cryptococcoma formation is predictive of high fungal burden outcome

Examination of infection dynamics over time led to the finding that cryptoccocal masses were often present before the final overwhelming infection and larval death (Fig. 3A). Indeed, a cryptococcoma was observed in every case, preceding infection by an average of 2 days (Fig. 3B) and was true for GFP, mCherry and mixed infection outcomes (Fig. 3C). Since initial inoculum was not predictive of the overwhelming infection, we decided to examine a potential role of cryptococcoma formation in influencing result, the predominance of GFP or mCherry positive cryptococcal cells in uncontrolled infection. Individual cryptococcomas were comprised of a single colour. Therefore, the colours of individual cryptococcomas were compared to the corresponding majority colour of overwhelming infection within individual fish. A clear relationship is demonstrated between each colour of cryptococcoma and the final outcome; a single (GFP or mCherry) cryptococcoma colour was significantly more likely to result in a single colour final outcome, with a corresponding finding for mixed cryptococcomas (Fig. 3D,E). Furthermore, the number of cryptococcomas within each fish increased the rate of fungal burden progression (Fig. 3F). These data demonstrate a key role of cryptococcoma formation during infection dynamics but do not address their role in dissemination.

**Figure 3.**
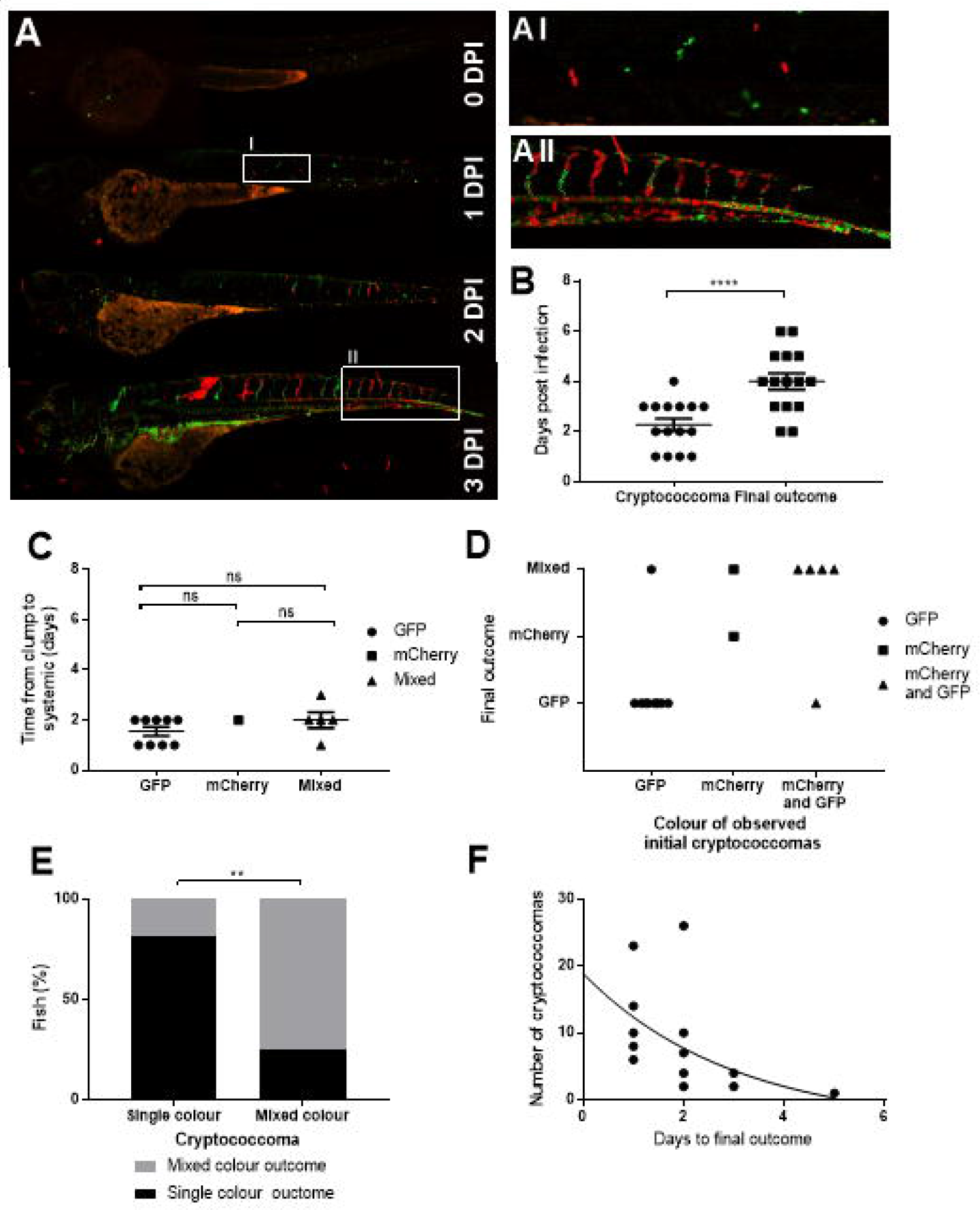
Cryptococcoma formation leads to uncontrolled infection. Infection of 2dpf AB larvae with 25cfu of a 5:1 ratio of GFP:mCherry KN99 *C. neoformans*. Larvae were imaged until 8dpf, or death (n=3, in each repeat 7, 10 and 12 larvae were used) **A.** Infection of AB wild-type larvae with 5:1 ratio of GFP:mCherry KN99 *C. neoformans*, at 0dpi, 1dpi, 2dpi and 3dpi **A I** Formation of cryptococcal masses at 1dpi **A II** Final infection outcome **B.** Time cryptococcoma first observed and time of final outcome observed (n=3, +/- SEM, Wilcoxon matched pairs test, ****p<0.0001) **C.** Time between cryptococcoma and final outcome grouped by final outcome majority colour (n=3, +/-SEM, One-way ANOVA, ns= not significant) **D.** Comparison of the colour (either GFP, mCherry or mixed) of *C. neoformans* in cryptococcomas, in relation to the final outcome majority *C. neoformans* colour **E.** Comparison of the colour of cryptococcomas, either single colour or mixed, with the colour of final outcome (n=3, **p<0.01, Fischer’s exact test) **F.** The number of cryptococcomas observed within individual larvae and how many days after observation final overwhelming infection was reached (n=3, non-linear regression, one-phase decay)

### Cryptococcal clonal expansion is more common in small blood vessels

It has been demonstrated that cryptococcal cells can become mechanically trapped in small blood vessels in the brain and subsequently disseminate (13). The mechanism by which cryptococci disseminate in this case is unknown but has been suggested to be via trancytosisis (13). We found that individual cryptococcal cells become trapped in the intersegmental vessels (ISVs) (Fig. 4A). We quantified the distribution of cryptococcomas and found that most (80.3%) were located in the smaller brain and trunk blood vessels (Fig. 4B). As cryptococcoma formation at the start of infection is seen in the smaller blood vessels, we determined whether clonal expansion was favoured in smaller blood vessels later in infection. We compared the ratio of GFP:mCherry between the trunk blood vessels and the caudal vein and found that in mixed infections there were single colour masses in the trunk vessels but dual colours in the larger caudal vein (Fig. 3AII; Fig. 4C), Thus, proliferation of cryptococci is favoured in small blood vessels.

**Figure 4.**
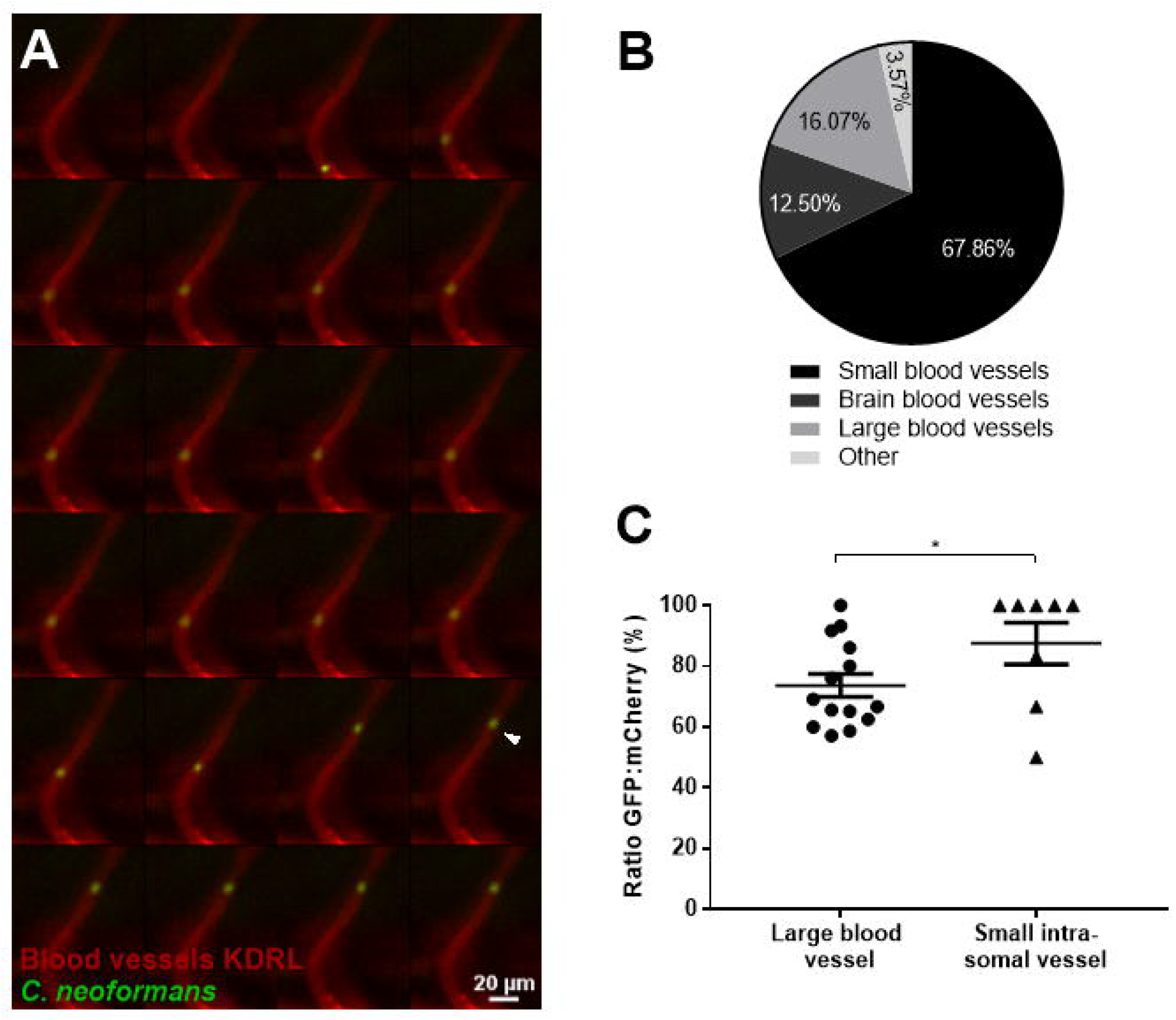
Cryptococcal cell trapping in small blood cells. **A.** Infection of KDRL mCherry blood marker transgenic line with 25cfu GFP *C. neoformans*, imaged immediately after infection. A single cryptococcal cell becomes trapped in the vasculature (white arrow), after moving from the bottom of the vessel to toward to top (left to right, time points 0.6 seconds) **B.** Infection of 2dpf AB larvae with 25cfu of a 5:1 ratio of GFP:mCherry KN99 *C. neoformans*. Larvae were imaged until 8dpf, or death (n=3, in each repeat 7, 10 and 12 larvae were used) Proportion of cryptococcomas observed in small inter-somal blood vessels, small brain blood vessels, large caudal vein or in other locations e.g. yolk, (n=3). **C.** The ratio of GFP:mCherry *C.neoformans* in the large caudal vein in comparison to the fifth inter-somal blood vessel, at uncontrolled infection time point (n=3, *p<0.05, +/-SEM, paired t-test).

### Clonal expansion results in vasodilation and disruption of vessel integrity leading to dissemination

We observed that enlargement of the cryptococcoma over time eventually led to invasion of the surrounding tissue at the site of infection (Fig. 1A). To determine how localised clonal expansion within the vasculature resulted in tissue invasion, we measured the width of blood vessels within the same infections, with and without cryptococcal growth. Blood vessels that contain cryptococcal cells were significantly larger than blood vessels that did not contain any cryptococcal cells at both 2hpi and 3dpi, although the magnitude of this difference was greater at 3dpi (Fig. 5A, B, SFig. 2), indicating cryptococcal growth within vessels throughout infection. The vessel width increase was proportional to the size of the cryptococcal mass inside the vessel at both 2hpi and 3dpi (Fig. 5C,D), although the size of cryptococcomas is larger at 3dpi. Injection of inert beads of a corresponding average cryptococcal cell size (4.5um) did not lead to formation of large masses, although there was a small but significant increase in vessel size at locations where beads did become trapped in the vasculature by 3dpi (Fig. 5E). Additionally, beads were observed stuck in the inter-segmental blood vessels significantly less frequently than live cryptococcal cells, with 13.6% of blood vessels containing beads compared to 89.0% containing cryptococcal cells. In addition, we imaged the small vessels of the brain and found that infected blood vessels were larger relative to blood vessels in the same location in control animals (Fig. 5F; Fig. 6G). This indicated that viable cryptococcal cells have an increased ability to become trapped and form masses compared to inert but similarly sized beads, and further implicates cryptococcal proliferation as a mechanism of inflicting vessel damage rather than a build-up caused by an initial blockage. Our finding that vessel diameter was enlarged suggested that blood vessels were vasodilating to reduce the total peripheral resistance to decrease blood pressure due to blood vessel blockage. Therefore, the role of localised clonal expansion in smaller blood vessels, the formation of cryptococcomas and their predictive capacity in terms of overwhelming infection, suggested a direct role in dissemination. We hypothesised that cryptococcomas were blocking vessels, increasing the force on the blood vessel walls, leading to vessel rupture and dissemination of cryptococci. To test our hypothesis we first established the relationship between tissue invasion and sites of clonal expansion within the vasculature. We found that in all cases tissue invasion occurred at sites of clonal expansion within the vasculature (19/19 tissue invasion events; Fig 6A). Furthermore, *C. neoformans* that had invaded the surrounding tissue were invariably the same colour (GFP or mCherry) as the vasculature cryptococcoma (Fisher’s exact test p<0.001, n=3, Fig. 6B). Ultimately, this observation indicates that clonal expansion is responsible for causing invasion of surrounding tissue, perhaps through vascular damage. To determine whether the vasculature was physically damaged sufficiently for cryptococcal cells to escape into the surrounding tissue, we examined blood vessels at high resolution at the sites of tissue invasion. We observed vessel damage and bursting at locations of cryptococcomas (Fig. 6C), in addition to tissue invasion events where the vasculature remained intact (Fig 6D), but we did not see this in non-infected vessels (SFig. S2). This suggested that dissemination was possible both via direct crossing of the endothelial barrier and by blood vessel disruption. Blood vessel integrity is maintained by individual cell integrity and the cell-cell junctions between vascular endothelial cells. To investigate vessel integrity, we generated a new zebrafish transgenic line expressing a fluorescently labelled reporter of the vascular endothelial cell junctional protein VE cadherin. Using this transgenic we investigated vessel integrity loss. We found that cryptococcal cells were located outside the blood vessel when vessels were either intact (Fig. 6E) or disrupted (Fig. 6F), in comparison to non-infected vessels (SFig. 3). This indicated that, in addition to transmigration across the vasculature, cryptococcal cells are able to disseminate via vessel damage.

**Figure 5.**
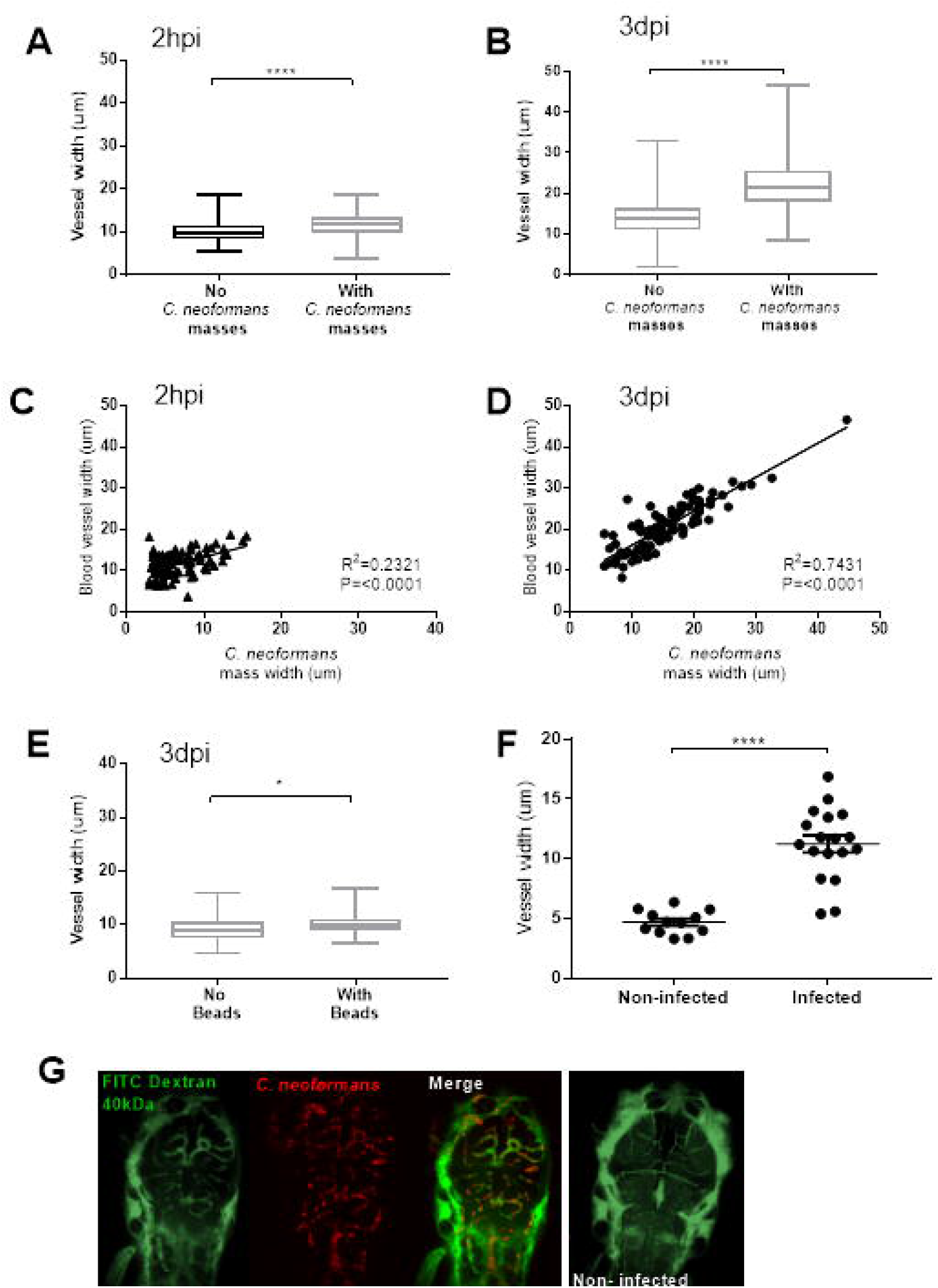
Localised clonal expansion proportionally increases vasculature size. **A-E** Infection of KDRL mCherry blood marker transgenic line with 1000cfu GFP *C. neoformans* or inert beads **A.** Vessel width with and without cryptococcal masses at 2hpi (n=3, +/- SEM, ****p<0.0001, unpaired t-test) **B.** Vessel width with and without cryptococcal masses at 3dpi (n=3, +/- SEM, ****p<0.0001, unpaired t-test) **C.** Relationship between *C. neoformans* mass and vessel width at 2hpi (n=3, linear regression) **D.** Relationship between *C. neoformans* mass and vessel width at 3dpi (n=3, linear regression) **E.** Vessel width with and without beads present at 3dpi (n=3, +/- SEM, *p<0.05, unpaired t-test). **F-G** Co-infection of mCherry *C. neoformans* with 40kDa FITC Dextran to mark blood vessels **F.** Comparison of infected brain vessels width to non-infected corresponding brain vessels (three infected fish analysed, +/- SEM, ****p<0.0001, paired t-test) **G.** Example image of infected and noninfected brain vessels.

**Figure 6.**
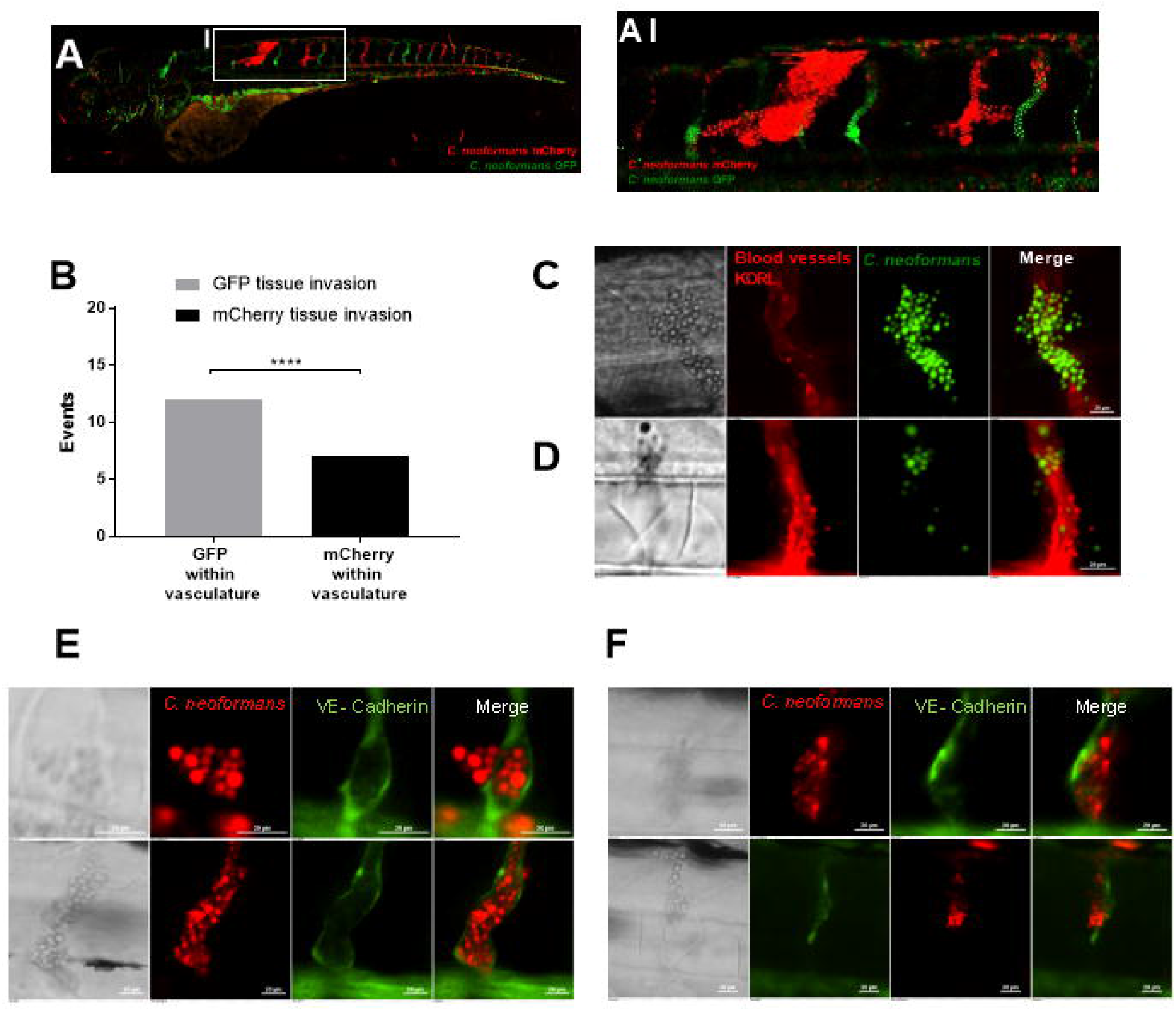
Dissemination events through vasculature damage. **A-D** Infection of KDRL mCherry blood marker transgenic line with 1000cfu GFP *C. neoformans* **A-AI** Example of dissemination of *C. neoformans* (mCherry) into the somite surrounding an existing mCherry cryptococcoma **B.** Comparison of colour of *C. neoformans* in the vasculature (GFP or mCherry), and the corresponding colour of dissemination events at the same location **C.** Dissemination from an intact blood vessel, with *C. neoformans* in the surrounding tissue suggested to be transcytosis **D.** Damaged blood vessels with *C. neoformans* in surround tissue **E-F** Infection of vascular-endothelium cadherin GFP tight junction (blood vessel marker) transgenic line with 1000cfu mCherry *C. neoformans* **E.** Intact tight junctions in the blood vessel endothelial layer, with *C. neoformans* in the surrounding tissue **F.** Damaged tight junctions in the blood vessel endothelial layer

## Discussion

Here, we describe and characterise a new mechanism for cryptococcal dissemination via proliferation with blood vessels, increasing local blood pressure and resulting in vessel rupture, disseminating cryptococci. *C. neoformans* invasion of the brain causes cryptococcal meningitis and meningoencephalitis. CNS infection with *C. neoformans* results in a high mortality rate, particularly in low and middle income countries. The mechanisms by which cryptococcal cells are able to cross into the brain during infection are therefore important in understanding disease progression and for the identification of potential therapeutic targets. Here we demonstrate a hitherto uncharacterised mechanism of dissemination from blood vessels, which occurs after cryptococcal trapping within small blood vessels and subsequent cryptococcal expansion causing vessel damage/bursting and invasion of the surrounding tissue by the cryptococcal cells.

Clinical studies and case reports indicate that vascular damage is caused by cryptococcal infection. Vascular damage during cryptococcosis may result from inflammation in the small blood vessels, seen predominantly in severe cases of cryptococcal meningitis (8–10,12). Indeed, in one retrospective study, infarcts were observed in as many as 30% of cryptococcal meningitis patients (12). Data collected in these studies suggest that infarcts occur during cryptococcal meningitis, perhaps through a mechanism of vasculitis resulting in vasculature damage. The physiological cause of infarcts in human cryptococcal infection has thus far not been identified. Our data suggest a possible mechanism whereby cryptococcoma formation results in vascular damage and cryptococcal dissemination associated with cortical infarcts.

We have shown that cryptococcal cells become trapped in the vasculature, proliferate, and disseminate into the surrounding tissue. Investigation of the integrity of the vasculature identified that damage resulted in cell-cell disruption. Vasculature damage was shown to be caused by cryptococcal proliferation, which physically pushes and damages the vasculature, typically where large cryptococcomas were observed within enlarged blood vessels.

Other fungal pathogens are known to cause infarcts and vasculitis in human infection (19,20). It is possible that a similar mechanism of localised expansion leading to vessel damage may occur following infection by multiple fungal pathogens. A case report of *Candida krusei* infection in a leg ulcer causing localised vasculitis (19) suggests that local fungal pathogen growth can damage the vasculature not only in the brain. In addition, infarcts, seen in cryptococcal meningitis, are also observed in meningitis caused by bacterial pathogens, for example *S. enterica* and *T. bacillus* (11,21).

In addition, we demonstrate that the presence and subsequent growth of cryptococcomas in the vasculature can lead to pathogen dissemination. This is consistent with post-mortem reports showing cryptococcal cells invading the brain, located next to brain capillaries that have a cryptococcal mass present (14). In cases where the vasculature is damaged, cryptococcal dissemination into surrounding tissue may be permitted by the ability of cryptococcal cells to escape the vasculature. We further suggest that vascular damage in cryptococcal infection is caused by cryptococcal trapping in small blood vessels and subsequent physical damage from localised clonal expansion, ultimately enabling spread from the vasculature to the brain.

Macrophages are an established intracellular niche in which cryptococcal cells are able to survive, replicate and escape (22). Diverse bacterial pathogens, for example *S. aureus, S. Enterica* and mycobacteria employ phagocytes as an intracellular niche (23–25). While most bacteria are degraded a small number successfully escape the phagocyte, this latter population continuing infection through clonal expansion, often leading to overwhelming infection (24). We investigated whether a population “bottle-neck” was present in cryptococcal infection. Through mixed infection of two fluorescent cryptococcal strains, we demonstrated that a high proportion of larvae have a final infection of mixed colour, indicating that multiple sources of cryptoccocal cells are responsible for high fungal burdens. Therefore it is not a single, and most likely not a small number of cryptococcal cells that is responsible for high fungal burden following a population bottleneck. How intracellular proliferation of cryptococcosis within host cells ultimately contributes to the progression is not known, although increased intracellular proliferation in macrophages is correlated with virulence in murine models and clinical isolates (26). Extracellular growth of cryptococcal cells is associated with infection in immunocompromised patients (4,5). In animal models of cryptococcal infection it is high number of extracellular fungal cells that is observed with high fungal burdens (4,5,16,27). We also tested what other factors might be associated with high fungal burden outcome. We determined that cryptococcoma formation and not initial inoculum was predictive of high fungal burden, suggesting that localised clonal expansion from distinct populations of cryptococcal cells, resulting in cryptococcoma formation, is responsible for uncontrolled infection.

Evidence that cryptococcal cells can become trapped in brain blood vessels was first adduced using a murine model. Furthermore, trapped cryptococcal cells appeared to transmigrate into the brain in real time (13). We observed cryptococcal trapping in small inter-segmental blood vessels in the zebrafish via a conserved trapping mechanism in similar sized blood vessels. *C. neoformans* trapping in small blood vessels may be urease dependent, particularly in organs such as the brain, but the mechanism is unknown (28). *C. neoformans* does not need to be able to proliferate in order to cross the BBB, as demonstrated using cryptococcal RAS1 and CDC42 mutants that can successfully transmigrate into the brain parenchyma (13). Although transmigration of non-proliferative *C. neoformans* across the BBB can occur, this does not negate a potential involvement of cryptococcal proliferation in crossing the BBB. We observed dissemination events resembling transmigration where the vasculature is intact, and also dissemination where the vasculature is damaged, where cryptococcal cells may escape into surrounding tissue, thus demonstrating a separate mechanism of dissemination. Infarcts are observed most often in severe cryptococcal meningitis (12), but it is not known if this is a cause or effect: it is possible that vascular damage occurs more frequently at a high fungal burden, or that high fungal burden arises due to due to a sudden introduction of extracellular yeast cells to the CNS. Additionally, dissemination events were observed at sites of localised cryptococcal expansion, where the fungal load is high. It is established that multiple mechanisms, including transmigration or via a macrophage Trojan horse, enable cryptococcal cells to cross the BBB (6,7). We observed dissemination via transcytosis at sites of cryptococcomas. This is suggestive that transcytosis events are promoted by the presence of large masses of cryptococcal cells, perhaps though increased interactions with the blood vessel wall. This implicates cryptococcal growth within blood vessels in facilitating dissemination events, not only through vasculature damage.

We have shown that localised clonal expansion is responsible for cryptococcoma formation and growth within the vasculature. A corresponding increase is vessel size was not observed for corresponding bead injections, consistent with growth and therefore increase in cryptococcoma size resulting in increased vasodilation due to vessel blockage. A direct effect of an increased size of vessels due to cryptococcal expansion has not been demonstrated previously and the increase in vessel size is likely to damage the integrity of the vessel wall. Damage to tight-junctions caused by *C. neoformans* has been suggested *in vitro*, in experiments where incubation of cryptoccocal cells with human brain microvascular endothelial cells caused damage of tight-junction transmembrane protein occludin (29). Here, using a VE-cadherin reporter, we demonstrate *in vivo* that at sites of cryptococcomas in the vasculature, cell-cell junctions are damaged resulting in a loss of vessel integrity and dissemination.

Our findings demonstrate a new mechanism of cryptococcal dissemination, whereby one or a small number of cryptococcal cells become trapped in the vasculature. Localised clonal expansion within the blood vessel leads to increased blood pressure resulting in failure of vessel integrity enabling cryptococcal dissemination, and suggesting that vessel damage caused by the presence of cryptococcoma in the vasculature may be responsible for vasculitis and infarcts observed in clinical cryptococcal meningitis cases.

## Methods and Methods

### Ethics statement

Animal work was carried out according to guidelines and legislation set out in UK law in the Animals (Scientific Procedures) Act 1986, under Project License PPL 40/3574 or P1A4A7A5E). Ethical approval was granted by the University of Sheffield Local Ethical Review Panel. Animal work completed in Singapore was completed under the Institutional Animal Care and Use Committee (IACUC) guidelines, under the A*STAR Biological Resource Centre (BRC) approved IACUC Protocol # 140977.

### Fish husbandry

Zebrafish strains were maintained according to standard protocols (30). Animals housed in the Bateson Centre aquaria at the University of Sheffield, adult fish were maintained on a 14:10-hour light/dark cycle at 28□ °C in UK Home Office approved facilities. For animals housed in IMCB, Singapore, adult fish were maintained on a 14:10-hour light/dark cycle at 28□°C in the IMCB zebrafish facility. We used the *AB* and *Nacre* strains as the wild-type larvae. The blood vessel marker *Tg(kdrl:mCherry)^S916^*, in addition to the vascular-cadherin marker line *TgBAC(ve-cad:GALFF)* (31), crossed to a *Tg(10xUAS:Teal)^uq13bh^* fluorescent transgenic zebrafish lines were used. *Tg(10xUAS:Teal)^uq13bh^* was generated by cloning Teal into the Gateway pME vector (pDON-221) using Gateway technology and the following primers:

- pME-Teal: 5’-**GGGGACAAGTTTGTACAAAAAAGCAGGCT**atggtga gcaagggcgaggag-3’ (gateway homology arm=bold)
- pME-Teal: 5’-**GGGGACCACTTTGTACAAGAAAGCTGGGTA**ctacttgt acagctcgtccatg-3’ (gateway homology arm=bold)

Subsequently a Gateway LR reaction was performed combining a p5E-10xUAS, pME-Teal and p5E-polyA placing the final 10xUAS:Teal and 10xUAS:Venus sequence into pDestTol2pA2AC (containing the α-crystallin promoter driving GFP in the zebrafish lens).

### *C. neoformans* culture

The *C. neoformans* variety *grubii* strain KN99, its GFP-expressing derivative KN99:GFP and mCherry-expressing derivative KN99:mCherry were used in this study. Culture were grown in 2 ml of yeast extract peptone dextrose (YPD) (all reagents are from Sigma-Aldrich, Poole, UK unless otherwise stated) inoculated from YPD agar plates and grown for 18 hours at 28 **°**C, rotating horizontally at 20 rpm. Cryptococcal cells were collected from 1ml of the culture, pelleted at 3300 g for 1 minute.

To count cryptococcal cells, the pellet was re-suspended in 1 ml PBS and cells were counted with a haemocytometer. Cryptococcal cells were pelleted again (3300g) and resuspended in autoclaved 10% Polyvinylpyrrolidinone (PVP), 0.5% Phenol Red in PBS (PVP is a polymer that increases the viscosity of the injection fluid and prevents settling of microbes in the injection needle), ready for micro-injection. The volume of PVP in Phenol red cryptococcal cells were re-suspended was calculated to give the required inoculum concertation.

### Zebrafish microinjection

An established zebrafish *C. neoformans* micro-injection protocol was followed (Bojarczuk et al., 2016). Zebrafish larvae were injected at 2 days post fertilisation (dpf) and monitored until a maximum of 10dpf. Larvae were anesthetised by immersion in 0.168 mg/mL tricaine in E3 and transferred onto 3% methyl cellulose in E3 for injection. 1nl of cryptococcal cells, where 1nl contained 25cfu, 200cfu or 1000cfu, was injected into the yolk sac circulation valley. For micro-injection of GFP fluorescent beads (Fluoresbrite® YG Carboxylate Microspheres 4.50μm). The bead stock solution was pelleted at 78g for 3 minutes, and re-suspended in PVP in phenol red as above for the required concentration. Micro-injection of 40kDa FITC-dextran (Sigma-Aldrich) at 3dpf in a 50:50 dilution in PVP in phenol red, injected 1nl into the duct of Cuvier. Larvae were transferred to fresh E3 to recover from anaesthetic. Any zebrafish injured by the needle/micro-injection, or where infection was not visually confirmed with the presence of Phenol Red, were removed from the procedure. Zebrafish were maintained at 28 °C.

### Microscopy of infected zebrafish

Larvae were anaesthetized 0.168 mg/mL tricaine in E3 and mounted in 0.8% low melting agarose onto glass bottom microwell dishes (MatTek P35G-1.5-14C). For low *C. neoformans* dose infection time points, confocal imaging was completed on a Zeiss LSM700 AxioObserver, with an EC Plan-Neofluar 10x/0.30 M27 lense. Three biological repeats contained 7, 10 and 12 infected zebrafish. Larvae were imaged in three positions to cover the entire larvae (head, trunk and tail) at 2 hours post infection, and at subsequent 24 hour intervals. After each imaging session, larvae were recovered into fresh E3 and returned to a 96-well plate.

A custom-build wide-field microscope was used for imaging transgenic zebrafish lines blood vessel integrity after infection with *C. neoformans*. Nikon Ti-E with a CFI Plan Apochromat λ 10X, N.A.0.45 objective lens, a custom built 500 μm Piezo Z-stage (Mad City Labs, Madison, WI, USA) and using Intensilight fluorescent illumination with ET/sputtered series fluorescent filters 49002 and 49008 (Chroma, Bellow Falls, VT, USA). Images were captured with Neo sCMOS, 2560 × 2160 Format, 16.6 mm x 14.0 mm Sensor Size, 6.5 μm pixel size camera (Andor, Belfast, UK) and NIS-Elements (Nikon, Richmond, UK). Settings for *Tg(kdrl:mCherry)* and *TgBAC(ve-cad:GALFF)* crossed to *Tg(10xUAS:Teal)^uq13bh^* GFP, filter 49002, 50 ms exposure, gain 4; mCherry, filter 49008, 50 ms exposure, gain 4. Settings for the GFP fluorescent beads were altered for GFP alone, filter 49002, 0.5 ms exposure, gain 4. In all cases a 50um z-stack section was imaged with 5um slices. Larvae were imaged at 2 hours post infection, and at subsequent 24 hour intervals. After each imaging session, larvae were recovered into fresh E3 and returned to a 96-well plate.

Co-injection of 40KDa FITC dextran with cryptococcal cells for imaging of vasculature in the brain was completed on 3dpf immediately after dextran injection, using a Ziess Z1 light sheet obtained using Zen software. A W-Plan-apochromat 20x/1. UV-Vis lense was used to obtain z-stack images using the 488nm and 561nm lasers and a LP560 dichroic beam splitter.

### Time-lapse microscopy of infected zebrafish

For time-lapse imaging of low *C.neoformans* dose infection, larvae were anaesthetised and mounted as described above, with the addition of E3 containing 0.168 mg/mL tricaine over-laid on top of the mounted *Nacre* larvae. Images were captured on the custom-build wide-field microscope (as above), with CFI Plan Apochromat λ 10X, N.A.0.45 objective lens, using the settings; GFP, filter 49002, 50 ms exposure, gain 4; mCherry, filter 49008, 50 ms exposure, gain 4. Images were acquired with no delay (~0.6 seconds) for 1 hour, starting <2mins after infection.

### Image analysis

Image analysis performed to measure the size of cryptococcal masses, and blood vessel width was completed using NIS elements. Fluorescence intensity of GFP and mCherry *C. neoformans* for low infection analysis was calculated using ImageJ software.

### Statistical analysis

Statistical analysis was performed as described in the results and figure legends. We used Graph Pad Prism 6 (v7.02) for statistical tests and plots.

## Acknowledgments

We thank Bateson Centre aquaria staff for their assistance with zebrafish husbandry. We thank Timothy Chico (University of Sheffield, UK) and Robert Wilkinson (University of Sheffield, UK) for help and advice on vascular biology.

JFG was supported by an A* Institute (Singapore) Doctoral Training Studentship. RJE was supported by a British Infection Association postdoctoral fellowship (https://www.britishinfection.org/). AKL was supported by a University of Queensland Postdoctoral Fellowship. BMH by an NHMRC/National Heart Foundation Career Development Fellowship (1083811). SAJ was supported by Medical Research Council and Department for International Development Career Development Award Fellowship MR/J009156/1 (http://www.mrc.ac.uk/). SAJ was additionally supported by a Krebs Institute Fellowship (http://krebsinstitute.group.shef.ac.uk/), and Medical Research Council Center grant (G0700091). SAR was supported by a Medical Research Council Programme Grant (MR/M004864/1) (http://www.mrc.ac.uk/). Light sheet microscopy was carried out in the Wolfson Light Microscopy Facility, supported by a BBSRC ALERT14 award for light-sheet microscopy (BB/M012522/1).

## Supplemental figure legends

**Figure S1**

**Injected ratio and number does not determine uncontrolled infection.** Infection of AB wild-type larvae with 5:1 ratio of GFP:mCherry KN99 *C. neoformans*, actual number of cryptococcal cells, both GFP and mCherry KN99 in 25cfu injected grouped by majority colour outcome

**Figure S2**

**Increased blood vessel width in cryptococcal infection.** Infection of KDRL mCherry blood marker transgenic line with 1000cfu GFP KN99 *C. neoformans* **A.** Example of vessels clear of cryptococcal cells blood vessels are intact **B.** Damaged blood vessels due to colonisation of *C. neoformans*

**Figure S3**

**Increased blood vessel width at sites of cryptococcomas.** Infection of vascular-endothelium cadherin GFP tight junction (blood vessel marker) transgenic line with 1000cfu mCherry *C. neoformans* **A.** *C. neoformans* within blood vasculature **B.** Example of vessels clear of cryptococcal cells tight junctions in the blood vessel endothelial layer are intact

